# Local changes in potassium ions modulate dendritic integration

**DOI:** 10.1101/2023.05.06.539205

**Authors:** Malthe Skytte Nordentoft, Athanasia Papoutsi, Naoya Takahashi, Mathias Spliid Heltberg, Mogens Høgh Jensen, Rune Nguyen Rasmussen

## Abstract

During neuronal activity the extracellular concentration of potassium ions ([K^+^]_o_) increases substantially above resting levels, but it remains unclear what role these [K^+^]_o_ changes play in dendritic integration of synaptic inputs. We used mathematical formulations and biophysical modeling to explore the role of activity-dependent K^+^ changes near dendritic segments of a visual cortex pyramidal neuron, receiving synaptic inputs tuned to stimulus orientation. We found that the fine-scale spatial arrangement of inputs dictates the magnitude of [K^+^]_o_ changes around the dendrites: Dendritic segments with similarly-tuned inputs can attain substantially higher [K^+^]_o_ increases than segments with diversely-tuned inputs. These [K^+^]_o_ elevations in turn increase dendritic excitability, leading to more robust and prolonged dendritic spikes. Ultimately, these local effects amplify the gain of neuronal input-output transformations, causing higher orientation-tuned somatic firing rates without compromising orientation selectivity. Our results suggest that local activity-dependent [K^+^]_o_ changes around dendrites may act as a “volume knob” that determines the impact of synaptic inputs on feature-tuned neuronal firing.

## Introduction

Throughout the nervous system, neuronal activity and ionic changes in the extracellular environment are bidirectionally linked. Despite this fact, extracellular ion changes are not traditionally considered an integral part of neuronal signaling and information processing. During neuronal activity, the concentration of several extracellular ionic species changes, but the most substantial relative change occurs for K^+^ ions. At rest, the extracellular concentration of K^+^ ([K^+^]_o_) in the brain is normally between 2.7 and 3.5 mM (Rasmussen et al., 2020; Somjen, 2002, 1979). During sensory stimulation, motor network activity, sleep oscillations, or behavioral state transitions, [K^+^]_o_ increases by 0.25–2 mM (Singer and Dieter Lux, 1975; Connors et al., 1979, 1982; Syková et al., 1974; Heinemann et al., 1990; Brocard et al., 2013; Amzica et al., 2002; Ding et al., 2016; Rasmussen et al., 2019), while it can rise to 7–12 mM during hypersynchronous neuronal activity (Heinemann et al., 1977; Poolos et al., 1987; Futamachi et al., 1974). The [K^+^]_o_ increase weakens the outward K^+^ driving force, resulting in a less negative K^+^ reversal potential 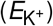, which powerfully affects the membrane potential (V_m_), excitability, and firing patterns of neurons (Malenka et al., 1981; Balestrino et al., 1986; Brocard et al., 2013; Hablitz and Lundervold, 1981; Rasmussen et al., 2019; Shih et al., 2013; Tong et al., 2014; Utzschneider et al., 1992; Rasmussen et al., 2017). Several sources contribute to [K^+^]_o_ changes but a major source is K^+^ efflux from excitatory glutamatergic synapses, in particular from NMDA receptors (Rice and Nicholson, 1990; Shih et al., 2013; Poolos et al., 1987). Previous work showed that such postsynaptic NMDA receptor-mediated K^+^ efflux can signal to presynaptic axons (Shih et al., 2013), pointing to local [K^+^]_o_ changes as a modulator of presynaptic transmission. However, whether local, postsynaptic, activity-mediated [K^+^]_o_ changes can modulate dendritic integration of synaptic inputs remains elusive.

Dendrites are the main receiving elements of neurons. They integrate information from thousands of presynaptic inputs made on dendritic spines and transmits it to the soma for generation of action potentials. An increasing body of literature has established that dendrites contain a variety of voltage-dependent ion channels which endow them with active electrical properties. For example, depolarization of the dendritic V_m_ activates these channels and can generate local action potentials, called dendritic spikes (Häusser and Mel, 2003; Major et al., 2013; Poirazi and Papoutsi, 2020; Stuart and Spruston, 2015). Dendritic spikes lead to nonlinear summation of synaptic inputs, which in turn boost the neurons’ input-output function and can enhance sensory feature selectivity (Smith et al., 2013; Lavzin et al., 2012; Bittner et al., 2017; Wilson et al., 2016). Interestingly, pharmacological manipulation of dendritic K^+^ currents affects dendritic excitability and dendritic spikes (Hoffman et al., 1997; Magee and Carruth, 1999). Thus, we here hypothesize that physiological synaptic activity-mediated changes in [K^+^]_o_ can locally regulate the active dendritic properties and shape nonlinear integration of inputs.

To test the hypothesis, we used mathematical formulations and biophysical modeling to look at the involvement of local K^+^ changes in the integration of synaptic inputs at the dendrite, with a particular focus on the spatial organization of synaptic inputs tuned to the stimulus orientation on dendritic branches of a pyramidal neuron in the visual cortex (Wilson et al., 2016). Here we show that the spatial arrangement of tuned inputs determines the magnitude of activity-dependent [K^+^]_o_ changes around dendrites. Specifically, dendritic segments with similarly-tuned synaptic inputs can attain substantially higher [K^+^]_o_ changes than segments with diversely-tuned inputs. These [K^+^]_o_ elevations in turn depolarize the 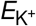 which increases dendritic excitability and prolongs dendritic spikes, but without compromising their stimulus selectivity. Ultimately, these local effects amplify the gain of neuronal input-output transformations, leading to higher firing rates at the soma without affecting feature selectivity. Our results suggest a prominent role for local activity-dependent K^+^ changes in shaping dendritic integration of synaptic inputs.

## Results

### Spatially clustered synaptic inputs cause local changes in [K^+^]_o_ and 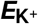 shifts

The spatial arrangement of synaptic inputs is an important factor for dendritic integration (Takahashi et al., 2012; Mel, 1993; Poirazi and Mel, 2001; Poirazi et al., 2003; Losonczy and Magee, 2006; Major et al., 2008; Weber et al., 2016). We hypothesized that dendritic segments with similarly-tuned inputs can attain higher [K^+^]_o_ and 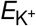 shifts than segments with diversely-tuned inputs, which could affect dendritic excitability (**Figure 1A**). For this we generated 10 µm segments populated with 7–13 synaptic spines, yielding spine densities comparable to neocortical tissue (Comer et al., 2020; Parker et al., 2020; Iascone et al., 2020). We chose this length scale because functional clusters, comprised of inputs with similar tuning preferences, typically are formed within 5–10 µm (Takahashi et al., 2012; Iacaruso et al., 2017; Scholl et al., 2017; Kerlin et al., 2019). To model feature-tuned synaptic inputs we used visual orientation tuning as a framework. Previous work identified dendritic segments with similar or diverse spine orientation tuning (Wilson et al., 2016; Jia et al., 2010). To capture this, we sampled spine orientation preferences from a circular normal distribution for the similarly-tuned regime and from a uniform distribution for the diversely-tuned regime (Wilson et al., 2016), and assigned tuning curves to individual spines (**Figure 1B** and **1C**). From the sampled segments, we obtained spine-averaged tuning curves and activity distributions (**Figure 1D** and **1E**). Using these we derived an activity factor for each orientation; signifying the ratio of expected spine activity in segments with similar versus diverse tuning preferences (**Figure 1F**; see Methods). This showed that for orientations close to the target orientation, here chosen as the soma preferred orientation (Δ target orientation: 0–20^◦^), spine activity levels are 2–5 times higher in segments with similarly-tuned inputs than in segments with diversely-tuned inputs. Conversely, for orientations far from the target orientation (Δ target orientation: 45–90^◦^), activity ceases almost entirely in segments with similar tuning while it stays constant for segments with diverse tuning, yielding activity factors below 1.

**Figure 1.**
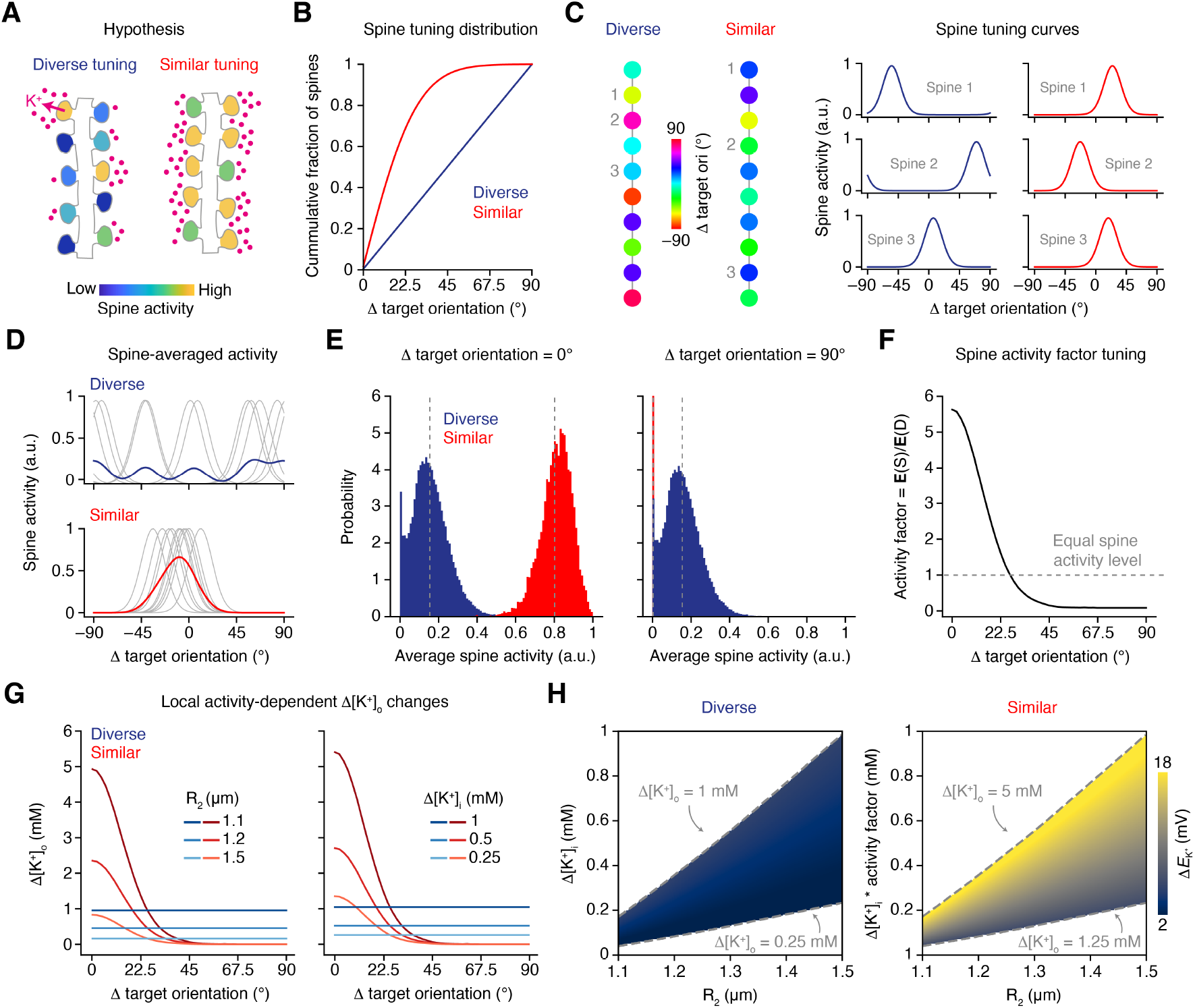
Spatially clustered synaptic inputs cause local changes in [K^+^]_o_ and 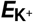 shifts. **(A)** Diagram of proposed hypothesis that dendritic segments with similarly-tuned inputs can attain higher [K^+^]_o_ increases than segments with diversely-tuned inputs. **(B)** Cumulative fraction of spine orientation preferences relative to target orientation (here chosen as preferred orientation of the soma) for dendritic segments with diverse or similar orientation tuning. Adopted from Wilson et al. (2016). **(C)** Example dendritic segments from the diverse and similar input tuning regimes (left). Tuning curves show orientation preference for the spines indicated on the corresponding dendritic segment (right). **(D)** Example spine-averaged tuning curves from dendritic segments from the diverse (top) and similar (bottom) tuning regimes. Tuning of individual spines are in gray and average is in color. **(E)** Probability distributions of average spine activity for the diverse and similar tuning regimes for the target (left) and orthogonal (right) orientations. Dotted lines indicate expectation value. **(F)** Dendritic spine activity factor as a function of stimulus orientation relative to target orientation. **(G)** Δ[K^+^]_o_ as a function of stimulus orientation relative to target orientation for the diverse and similar tuning regimes. Left: different extracellular space radii (R_2_) with constant [K^+^]_i_ reductions (Δ[K^+^]_i_ = 1 mM for diversely-tuned regime). Right: different Δ[K^+^]_i_ with constant R_2_ (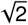). **(H)** Heat-maps showing range of 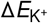 for the diverse (left) and similar (right) tuning regimes as a function of Δ[K^+^]_i_ and R_2_ for the target orientation. Dotted lines indicate the corresponding Δ[K^+^]_o_ range.

We then asked how the spine activity patterns might manifest in [K^+^]_o_ and 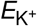 changes. To explore this we assumed the following conditions were true: 1) Local increases in [K^+^]_o_ are proportional to spine activity level; 2) Local increases in [K^+^]_o_ are caused by K^+^ efflux from the intracellular space; and 3) K^+^ ions in the extracellular space, for the spatial and temporal scale considered here (∼10 µm and ∼300 ms after synaptic activity onset, respectively; see Methods), can be described as well-mixed due to the high extracellular K^+^ free diffusion rate 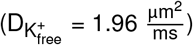 (Ellingsrud et al., 2020; Chen and Nicholson, 2000), meaning that the [K^+^]_o_ gradient along the 10-µm segment is zero in space and time. We can thus approximate [K^+^]_o_ changes in the vicinity of dendritic segments by multiplying the intracellular K^+^ concentration ([K^+^]_i_) change by the volume fraction between the intra- and extracellular space:

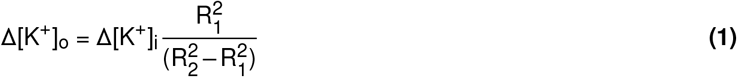

Here Δ[K^+^]_o_ and Δ[K^+^]_i_ are the changes in [K^+^]_o_ and [K^+^]_i_, respectively. The dendritic segment is considered a cylinder with radius R_1_, encapsulated by the extracellular space, also described by a cylinder with radius R_2_. By keeping the dendritic thickness constant (R_1_ = 1 µm) (Yamada and Kuba, 2021), we limit our parameter space to two parameters: Δ[K^+^]_i_ and R_2_. Although we do not have data on dendritic [K^+^]_i_ changes nor do we know the exact extracellular space size in the vicinity of dendritic segments, we can use equation 1 to estimate [K^+^]_o_ changes by varying Δ[K^+^]_i_ and R_2_ within realistic ranges (Grafe et al., 1982; Tønnesen et al., 2018; Pallotto et al., 2015). By multiplying Δ[K^+^]_i_ for the diversely-tuned input regime with the activity factor (**Figure 1F**), we obtain estimates of [K^+^]_o_ changes near segments with similarly-tuned inputs. Notably, for orientations close to the target orientation (Δ target orientation: 0–20^◦^), [K^+^]_o_ can increase by 0.2–4 mM more in segments with similar tuning preferences than in segments with diverse tuning, depending on stimulus orientation, Δ[K^+^]_i_ and extracellular space size (**Figure 1G**). Conversely, for orientations far from the target (Δθ^◦^ from target: 45–90^◦^), [K^+^]_o_ increases are 0.2–1 mM higher in segments with diversely-tuned inputs.

Next we constrained the Δ[K^+^]_o_ range by considering values of 0.25–1 mM, translating to depolarizing 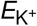 shifts of 1.6 to 5.9 mV (from a baseline of –80 mV), in the diversely-tuned regime realistic and conservative. This was justified by *in vivo* [K^+^]_o_ measurements using microelectrodes (Malenka et al., 1981; Singer and Dieter Lux, 1975; Connors et al., 1979; Syková et al., 1974; Somjen et al., 1976; Heinemann et al., 1990; Amzica et al., 2002; Ding et al., 2016; Rasmussen et al., 2019); a technique which averages in space and time and likely underestimates true local [K^+^]_o_ changes (Nicholson, 1993; Fröhlich et al., 2008; Frant and Ross, 1970; Walz and Hertz, 1983). Using this Δ[K^+^]_o_ interval we determined the corresponding Δ[K^+^]_i_ and R_2_ values to create a realistic parameter set, and obtained Δ[K^+^]_o_ values within 1.25–5 mM for the similarly-tuned input regime when stimulated with the target orientation (**Figure 1H**). This translates to depolarizing 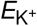 shifts of 6 to 18 mV, which could have significant effects on dendritic electrical properties. Together, these data suggest that dendritic segments receiving synaptic inputs with similar tuning preferences can attain substantially higher [K^+^]_o_ and 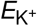 changes compared with segments receiving diversely-tuned inputs.

### 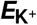 shifts regulate active dendritic properties

We then asked what implications the 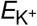 changes might have on dendritic synaptic integration. Here we focused on dendritic segments with similarly-tuned inputs. This was motivated by: 1) We only saw orientation-dependent and large 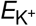 changes in segments with similar tuning preferences (**Figure 1G** and **1H**); and 2) Previous work have shown that spatial clustering of co-active inputs is an important factor for dendritic spike initiation (Takahashi et al., 2012; Mel, 1993; Poirazi and Mel, 2001; Poirazi et al., 2003; Losonczy and Magee, 2006; Major et al., 2008; Weber et al., 2016). For this analysis we established a biophysical model to simulate the V_m_ of a dendritic segment during synaptic input stimulation, referred to as a “point-dendrite” model due to its similarities with a conventional point-neuron model (Dayan and Abbott, 2001) (**Figure 2A**; see Methods). To mimic the active properties of dendrites, the model included an array of intrinsic ion channels as well as AMPA and NMDA receptors (Hay et al., 2011; Park et al., 2019; Shai et al., 2015) (**Table S1**). The number of spines and their orientation tuning was sampled as in **Figure 1** to resemble a dendritic segment receiving inputs with similar tuning. Synaptic inputs were simulated by delivering three stimulation trains in which AMPA and NMDA receptors were activated with a peak current depending on the spine’s orientation tuning and with a Poisson-distributed delay between spine activation (mean = 30 ms). From this we obtained V_m_ dynamics that reflected the stimulus orientation: For orientations close to the target orientation, we observed regenerative activation of voltage-gated sodium, calcium, and NMDA conductances, leading to dendritic spikes (mixture of sodium, calcium, and NMDA spikes), while for the orthogonal orientation the V_m_ did not change (**Figure 2B**). Next we tested the effect of 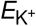 shifts (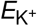 ∈ [0–18] mV) on dendritic spike properties (see Methods). Shifting 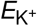 caused a linear broadening of dendritic spikes (**Figure 2C** and **2D**); for example, for a 12 mV 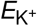 shift the duration of dendritic spikes increased by 60% compared to no 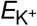 change. Furthermore, the amount of excitatory synaptic drive needed to transition the dendritic V_m_ from subthreshold input summation to suprathreshold spiking decreased substantially with increasing 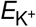 shifts (**Figure 2E** and **2F**).

**Figure 2.**
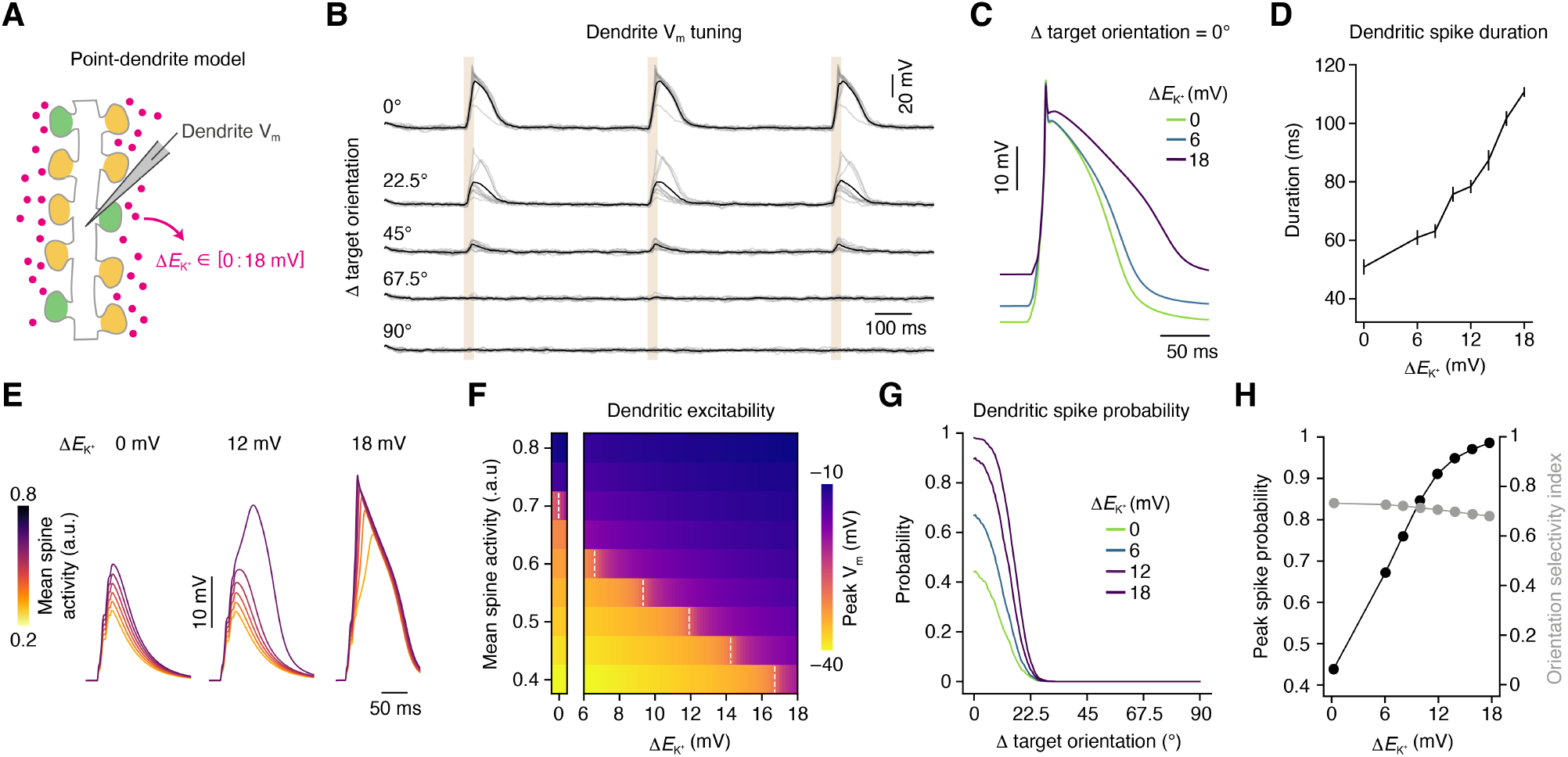
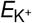 shifts regulate active dendritic properties. **(A)** Diagram of the biophysical point-dendrite model. To test the effect of local [K^+^]_o_ elevations we imposed 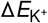 shifts within the interval of 0-18 mV. **(B)** Example dendrite V_m_ traces for the similar input tuning regime as a function of stimulus orientation. Individual trials are in gray and average is in black. Shaded regions indicate synaptic stimulation timing. **(C)** Example V_m_ traces highlighting dendritic spike duration (time above –30 mV) as a function of 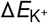 at the target orientation. **(D** Dendritic spike duration as a function of 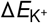 at the target orientation. Error bars are mean ± s.e.m. (*N* = 10 simulations). **(E)** Example traces highlighting V_m_ response as a function of 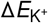 and synaptic drive. **(F)** Heat map showing peak V_m_ depolarisation as a function of 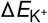 and synaptic drive. Dotted lines indicate the transition to dendritic spiking. **(G)** Dendritic spike probability as a function of 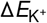 and stimulus orientation relative to target orientation. Note that 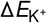 values denote changes at the target orientation. **(H)** Dendritic spike probability at the target orientation (left axis) and orientation selectivity index (right axis) as a function of 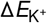. See also **Figure S1** and **Table S1**.

In our model, three main parameters control dendritic spike generation: the number of dendritic spines (N), mean spine activity (w), and 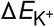. By simulating different parameter configurations with dendritic spike occurrence as binary output measure, we derived the function which describes the minimum 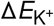 needed to trigger dendritic spiking given N and w (**Figure S1**):

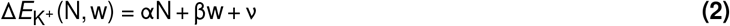

Using equation (2) we estimated the probability of generating a dendritic spike for a given stimulus orientation and 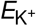 shift magnitude (**Figure 2G**). The chance of eliciting dendritic spikes rose drastically as a function of increasing 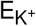 shifts (**Figure 2G** and **2F**); for example, the probability of spiking to the target orientation increased from 45% to 90% when shifting the 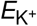 by 12 mV. Interestingly, the dendritic spiking selectivity was largely constant as the probability of spiking also increased for orientations away from the target orientation (**Figure 2G** and **2H**; Orientation selectivity index: 0.74, 0.73, and 0.69 for 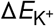 = 0, 6, and 18 mV, respectively). Altogether, these results show that local K^+^ changes near dendritic segments with similarly-tuned synaptic inputs can increase excitability, prolong dendritic spikes, and boost the robustness of generating dendritic spikes without compromising their feature selectivity.

### 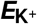 shift alters the current-voltage attractor landscape in dendrites

To understand the general biophysical principles governing the regulation of dendritic spikes by local K^+^ changes we turned to dynamical systems theory (Strogatz, 2015). Dendritic spiking can be described by the instantaneous attractor landscape of current-voltage (I-V) relations (Major et al., 2013), and we therefore generated such landscapes using our point-dendrite model. With only the intrinsic ion channels active, the system is down-stable and the V_m_ is attracted to a single hyperpolarized fixed point (**Figure 3A**). Including NMDA receptors changes the system to bistable, and a large attractor basin together with a depolarized fixed point, corresponding to dendritic spiking, is created. The current barrier preventing the system from transitioning to an up-stable state, where V_m_ is only attracted to the depolarized fixed point, is overcome by the activation of AMPA receptors. As the activity of AMPA and NMDA receptors wears off, in conjunction with high activity of K^+^ channels, the system is again pushed back to the down-stable state.

**Figure 3.**
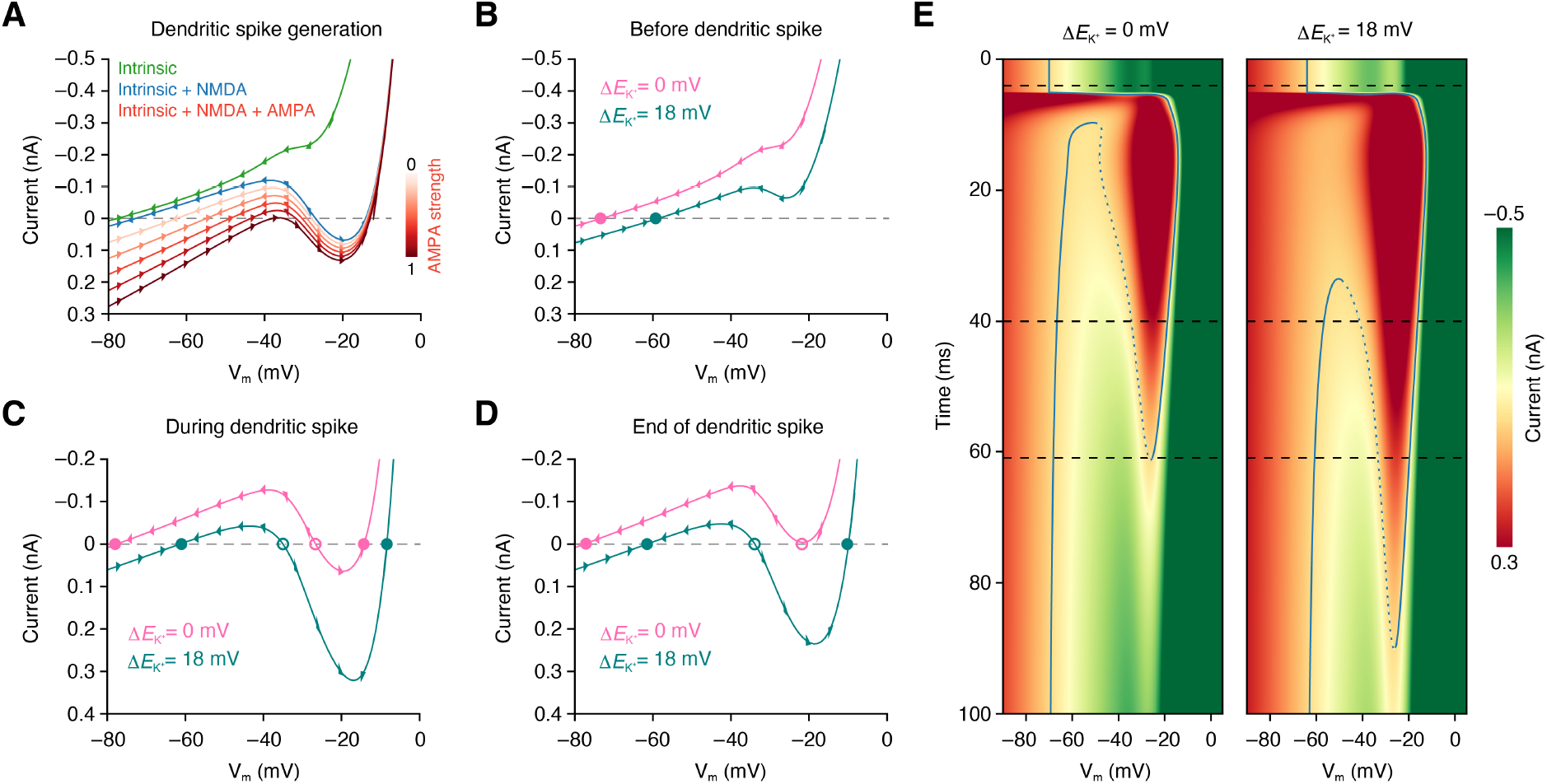
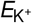 shift alters the current-voltage attractor landscape in dendrites. **(A)** I-V curves during the generation of dendritic NMDA spike. Green: down-stable state with only intrinsic ion channels active. Blue: bistable state with intrinsic ion channels and NMDA receptor active. Red: Bi- and up-stable state with intrinsic ion channels and NMDA and AMPA receptors active. Arrows indicate system flow direction. **(B)** Down-stable state before dendritic spike generation without and with 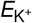 shift. Solid points indicate stable fixed points. (**C, D**) Bistable states during and at the end of dendritic spike without and with 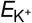 shift. **C** shows the hypothetical system with peak NMDA receptor conductance, without AMPA receptor activation, right before spike initiation, and **D** shows the system when outward and inward currents match for the system without 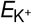 shift around the end of the spike. Solid and open points indicate stable and unstable fixed points, respectively. (**E**) Heat maps showing the temporal evolution of the I-V landscape without and with 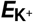 shift. The corresponding I-V curves shown in **B, C** and **D** are indicated with black dotted lines, and full and dotted blue lines indicate stable and unstable fixed points, respectively. See also **Movie S1**.

Shifting the 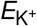 alters the I-V curve attractor landscape in three fundamental ways (**Figure 3B–E**; **Movie S1**). First, for the down-stable state, it moves the fixed point to a more depolarized V_m_, as well as reducing the net outward current across all V_m_ levels (**Figure 3B**). Second, for the bistable state, it lowers the current barrier needed to transition the system to the up-stable state, as well as deepens the attractor basin (**Figure 3C**). Finally, by reducing the net outward currents at depolarized V_m_ levels, it prolongs the duration of the up- and bistable states (**Figure 3C–E**). These dynamical system properties are all explained by the weakened K^+^ driving force through K^+^ channels, and fully predict the 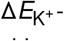 -mediated effects we observed on dendritic spiking (**Figure 2**). It should be noted that while the results presented here focus on the NMDA spike, the key observations should transfer to sodium and calcium spikes since these also rely on V_m_ depolarization to activate their active conductances. Collectively, this analysis demonstrates that the effects of local K^+^ changes on synaptic integration are underlied and can be predicted by altered dynamical system properties of dendritic spikes.

### Local dendritic 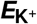 shifts induces neuronal firing gain modulation

Motivated by the intriguing finding that local [K^+^]_o_ increases near dendritic segments with similarly-tuned inputs can boost dendritic spike generation without compromising feature selectivity (**Figure 2G** and **2H**), we in our final investigations asked what functional consequences this might have on neuronal output. For this we constructed a neuron model using a fractal tree, mimicking the generic compartmentalization and fractal dimensionality of the apical dendritic tree of pyramidal neurons (Smith et al., 2021; Porter et al., 1991) (**Figure 4A**). To evoke somatic firing we stimulated distal dendrites with orientation-tuned excitatory inputs while shifting 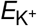 values in segments with diverse or similar tuning preferences (1:1 ratio of segment types) in three conditions: no change, small changes, or large changes, based on the [K^+^]_o_ changes determined in **Figure 1** (**Figure 4A** and **4B**; see Methods). This revealed that somatic firing rates were significantly amplified when 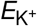 values were shifted locally (**Figure 4C** and **4D**; *P* = 10^−16^ and 10^−30^ for small and large 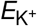 shifts, respectively, one-tailed Student’s *t* -test, *N* = 25 simulations). We determined that this form of gain modulation was remarkably consistent with a multiplicative transformation (**Figure S2**; gain coefficient = 1.24 and 1.96 for small and large 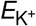 shifts, respectively), similarly to previously shown in the visual cortex (Polack et al., 2013). Importantly, and in congruence with what we observed in the dendrites (**Figure 2H**), the soma firing selectivity was not changed by 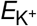 shifts despite increases in firing rate also for orientations away from the target orientation (**Figure 4C** and **4D**; orientation selectivity index: 0.89, 0.89, and 0.87 for no, small, and large 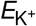 shifts, respectively; *P* = 0.07, χ^2^ fit, *N* = 25 simulations). As expected, the gain effect became stronger or weaker when shifting the relative fraction of dendritic segments towards more segments with similarly- or diversely-tuned inputs, respectively (**Figure S3**). To elucidate how dendritic 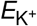 shifts cause dendritic gain modulation, we measured the area under the curve of the integrated V_m_ signal arriving at the base of the dendritic trunk during a synaptic stimulation burst. To exclude back-propagating action potentials (bAPs), we here turned off the voltage-gated sodium channels in the soma and trunk. This showed that the strength of the integrated dendritic signal, received by the soma, rose substantially with increasing 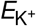 shifts (**Figure 4E**); for example, the signal strength at the target orientation increased by around 33% when imposing large 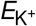 shifts compared to no 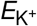 shifts.

**Figure 4.**
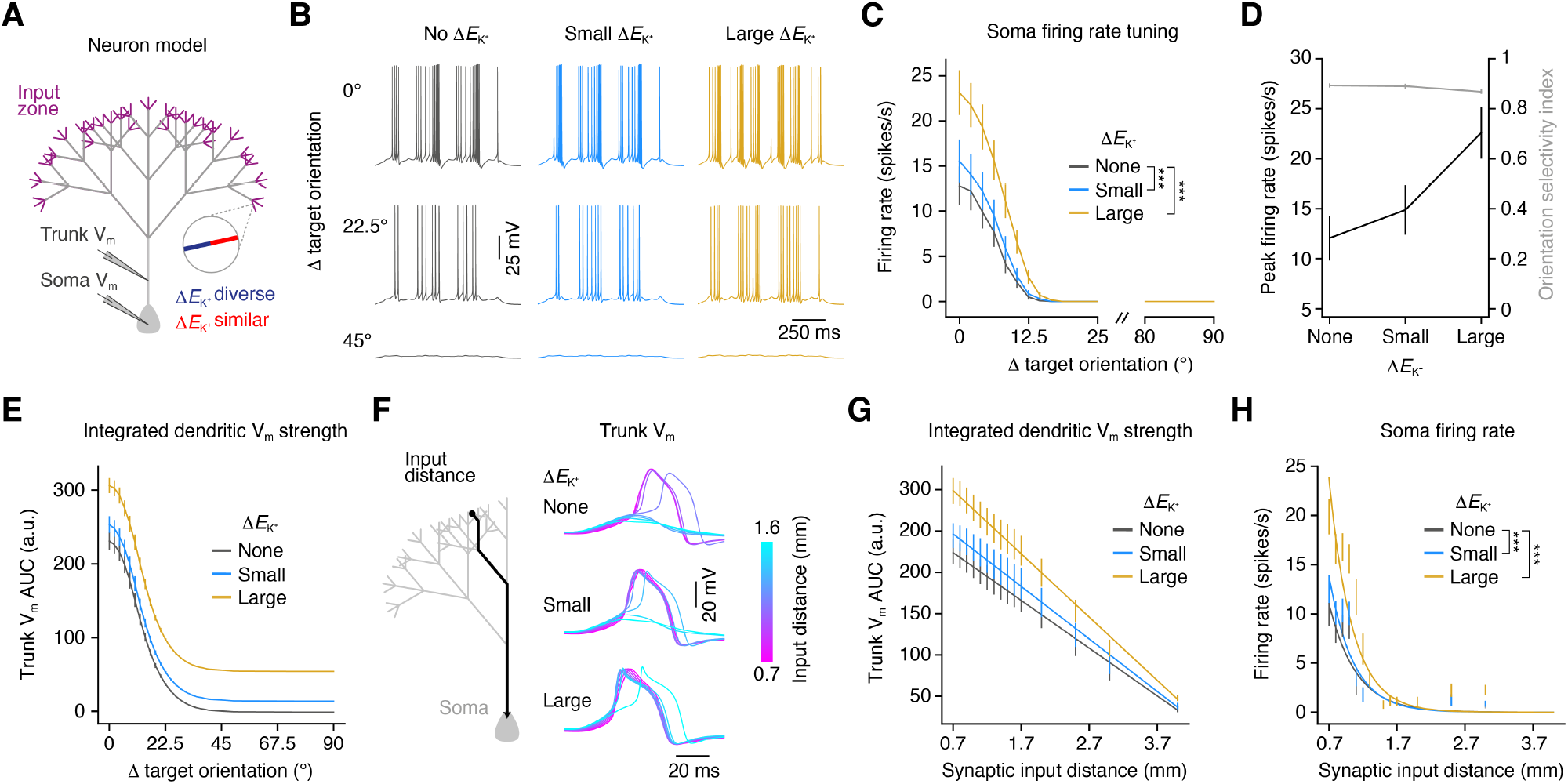
Local dendritic 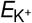 shifts induces neuronal firing gain modulation. **(A)** Diagram of the neuron model. The neuron received orientation-tuned synaptic input at the distal dendrites, and 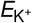 shifts corresponding to segments with diverse or similar tuning preferences were imposed across the dendritic tree. **(B)** Example soma V_m_ traces obtained as a function of dendritic 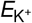 shift magnitude and stimulus orientation. **(C)** Soma firing rate tuning curve as a function of dendritic 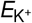 shift magnitude. Error bars are mean±s.e.m. (*N* = 25 simulations). \*\*\**P* = 10^−16^ and 10^−30^ for small and large 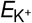 shifts, respectively, one-tailed Student’s *t* -test. **(D)** Soma peak firing rate at target orientation and orientation selectivity index as a function of dendritic 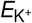 shift magnitude. **(E)** Area under the curve for the basal trunk V_m_ measurements as a function of stimulus orientation relative to target orientation and dendritic 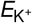 shift magnitude. Error bars are mean ± s.e.m. (*N* = 25 simulations). Note that bAPs were here abolished by silencing voltage-gated sodium channels in the trunk and soma. **(F)** Example basal trunk V_m_ traces obtained at the target orientation as a function of synaptic input distance to soma and dendritic 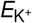 shift magnitude. **(G)** Area under the curve for the basal trunk V_m_ measurements at the target orientation as a function of synaptic input distance to soma and 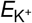 shift magnitude. Error bars are mean ± s.e.m. (*N* = 25 simulations) and solid lines represent linear fit. **(H)** Soma firing rate at the target orientation as a function of synaptic input distance to soma and 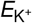 shift magnitude. Error bars are mean±s.e.m. (*N* = 25 simulations) and solid lines represent simple exponential fit. \*\*\**P* = 10^−22^ and 10^−46^ for small and large 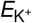 shifts, respectively, one-tailed Student’s *t* -test. See also **Figures S2** and **S3** and **Table S2**.

Finally, we asked what functional benefits there might be to this K^+^-mediated gain modulation. A major task of dendrites is to transmit incoming inputs to the soma for output generation. This is an inherent challenge, especially for distal inputs to large neurons, as the voltage signal tends to attenuate as a function of distance traveled and branching points (Gooch et al., 2022; Stuart and Spruston, 1998). We therefore speculated that one function of K^+^-mediated gain modulation might be to promote neurons transmitting signals over larger dendritic distances. To test this, we manipulated the synaptic input distance to the soma by varying the dendritic trunk length while measuring the integrated V_m_ signal at the base of the trunk or soma firing rate (**Figure 4F-H**). Interestingly, due to the local gain of the synaptic input, the depolarizing voltage plateau is able to travel a substantially longer distance in the 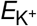 -shifted regime before reaching the level of the non-shifted regime; for example, the integrated V_m_ signal for a neuron with inputs 1700 µm away from the soma and large 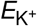 shifts is comparable to a neuron with inputs 700 µm away and no 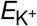 shifts, yielding a 140% increase in propagation distance (**Figure 4F** and **4G**). These effects translated to increases in soma firing rates for a given synaptic input distance when dendritic 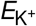 values were shifted (**Figure 4H**; *P* = 10^−22^ and 10^−46^ for small and large 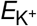 shifts, respectively, one-tailed Student’s *t* -test, *N* = 25 simulations). Together, these results show that local activity-dependent [K^+^]_o_ increases and 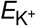 shifts in dendrites increase the gain of feature-tuned firing of neurons without comprising feature selectivity.

## Discussion

We have developed mathematical formulations and biophysical models to address an open question in neuroscience: How do local activity-dependent changes in [K^+^]_o_ affect dendritic integration of sensory-tuned synaptic inputs? Our work provides three major insights to this fundamental question. First, the fine-scale spatial arrangement of orientation-tuned synaptic inputs determines the magnitude of activity-dependent [K^+^]_o_ changes locally around dendritic segments; that is, segments with similarly-tuned inputs can attain substantially higher [K^+^]_o_ increases than segments with diverse inputs. Second, these [K^+^]_o_ elevations in turn depolarize the 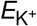 which enhances dendritic excitability, causing more robust and longer-lasting sensory-evoked dendritic spikes. Finally, the local dendritic effects promote gain amplification of neuronal input-output functions, resulting in increased somatic responsiveness without affecting feature selectivity of the neuron. Our results, therefore, suggest a prominent and previously overlooked role for local activity-dependent changes in K^+^ concentration in regulating dendritic computations, shedding new light on the mechanisms underlying sensory integration in neurons.

Dendritic processing of synaptic inputs depends on the spatial and temporal organization of the inputs: spatially dispersed and asynchronous inputs are summed linearly, while inputs that are spatially close and synchronized are summed nonlinearly and can facilitate the generation of dendritic spikes (Takahashi et al., 2012; Mel, 1993; Poirazi and Mel, 2001; Poirazi et al., 2003; Gasparini et al., 2004; Losonczy and Magee, 2006; Major et al., 2008; Weber et al., 2016). Here we propose the novel idea that another important function of grouping co-tuned synaptic inputs close in space is to generate higher activity-dependent [K^+^]_o_ increases, creating local and feature-tuned dendritic “[K^+^]_o_ hotspots.” Such [K^+^]_o_ hotspots, potentially attaining up to 5 mM K^+^ increases relative to baseline (**Figure 1**), have the capacity to markedly affect local dendritic processing by dampening K^+^ currents. For example, we here show that it can reduce the amount of excitatory drive needed to trigger dendritic spiking, as well as broadening the dendritic spikes (**Figures 2** and **3**). This is in congruence with previous work showing that K^+^ channels are highly expressed in dendrites (Johnston et al., 2000), and pharmacological or genetic disruption of dendritic K^+^ channels (Hoffman et al., 1997; Magee and Carruth, 1999; Sun et al., 2011; Johnston et al., 2000), or global [K^+^]_o_ elevations (Gasparini et al., 2004), increases the excitability of dendrites and prolongs dendritic spikes. Such broadening of dendritic spikes could potentially enhance the capacity of the neuron to integrate temporally delayed excitatory inputs (Du et al., 2017) and might be a cellular mechanism involved in short-term memory.

While we here only focused on the postsynaptic impact of local [K^+^]_o_ changes, it is reasonable to predict that activity-dependent [K^+^]_o_ hotspots could also affect the presynaptic terminals (Malenka et al., 1981). Increases in [K^+^]_o_ depolarize axons, which can broaden action potentials and increase presynaptic calcium entry (Geiger and Jonas, 2000; Sasaki et al., 2011), leading to enhanced glutamate release and stronger synaptic transmission. Such amplification of local synaptic integration, through pre- and postsynaptic mechanisms, could play important roles in neuronal circuit development and long-term potentiation by supporting spike-timing-dependent plasticity (Caporale and Dan, 2008; Feldman, 2012). Moreover, we did not assess GABAergic inhibition, known to play important roles for regulating dendritic activity (D’Aquin et al., 2022; Chiu et al., 2013). Interestingly, physiological elevations of [K^+^]_o_ can depolarize the reversal potential of GABA_A_ receptors through reduced activity of the potassium-chloride co-transporter (Thompson and Gahwiler, 1989; Jensen et al., 1993; Kaila et al., 1997), which in turn dampens inhibitory strength or even transforms GABAergic signaling to excitatory (Alfonsa et al., 2023; Kaila et al., 2014). We thus speculate if local [K^+^]_o_ increases near dendritic segments with similarly-tuned excitatory inputs are sufficiently large to cause disinhibition by reducing the inward driving force for chloride ions. Such feature-tuned local disinhibition could increase neuronal feature selectivity or create a temporal window for dendritic plasticity. To obtain a more complete picture, future models should aim to incorporate [K^+^]_o_-mediated regulation of excitatory presynaptic activity and GABAergic transmission.

It remains an open question what the functional role of the [K^+^]_o_-dependent dendritic integration modulation described here might be. We show that the combined effect of local activity-dependent [K^+^]_o_ increases is to cause multiplicative gain modulation of the neuronal input-output function by boosting the effectiveness of orientation-tuned synaptic distal inputs (**Figure 4**). This makes the neuron more responsive and could contribute to increasing the signal-to-noise ratio of visual cortical neurons while, importantly, maintaining their orientation selectivity. Furthermore, the dendritic response to inputs arriving on the apical dendrites have been implicated in context-dependent sensory processing and consciousness (Fişek et al., 2023; Takahashi et al., 2016; Suzuki and Larkum, 2020). Fascinatingly, dendritic spikes in apical dendrites of layer 5 pyramidal neurons are causally linked to the perceptual detection threshold in awake mice (Takahashi et al., 2016). Moreover, a preliminary report suggests that densely localized thalamic inputs to apical dendrites might gate sensory perception by facilitating dendritic spikes (Guest et al., 2021). It is thus tempting to speculate that spatial clustering of similarly-tuned thalamic inputs to apical dendrites promotes dendritic spikes, and lowers the perceptual threshold, by means of local activity-dependent [K^+^]_o_ increases. In this context, it is important to note that while we here chose to use visual orientation tuning as a framework, our proposed mechanistic concept is agnostic to sensory feature, brain region, or animal species. The fundamental requirement for the emergence of dendritic [K^+^]_o_ hotspots is that synaptic inputs are activated on a spatially and temporally synchronized scale. Hence, this mechanism could play a role in dendritic computations in diverse brain regions such as the hippocampus, motor cortex, visual cortex, and somatosensory cortex where spatiotemporally synchronized inputs have been observed (Takahashi et al., 2012; Iacaruso et al., 2017; Scholl et al., 2017; Kerlin et al., 2019; Wilson et al., 2016; Winnubst et al., 2015).

Our approach also comes with its limitations. One is that we did not model the temporal kinetics of the local [K^+^]_o_ dynamics. Our mean-field approach quantifies average [K^+^]_o_ changes after synaptic activation and thus lacks temporal resolution. We chose this approach because several important parameters have yet to be experimentally determined, including the voltage-dependent fraction of K^+^ currents of total NMDA currents and the precise geometry of the extracellular space surrounding functionally-mapped dendritic spines, which prevents precise simulations of time-dependent [K^+^]_o_ dynamics. We here assumed that the system goes asymptotically towards the well-mixed state following synaptic activation, and we estimate that this is reached ∼250–300 ms after input onset, but we cannot infer the nature of the spatiotemporal [K^+^]_o_ dynamics before this time point. More experimental knowledge about these parameters would enable us to make a full-scale electrodiffussive model of the extracellular space, improving the temporal resolution as well as rendering the end-plate boundary conditions superfluous. Additionally, although our work predicts the existence of local activity-dependent [K^+^]_o_ hotspots, this remains to be experimentally tested. The gold-standard for measuring [K^+^]_o_ dynamics in the brain is with K^+^-selective microelectrodes (Singer and Dieter Lux, 1975; Connors et al., 1979, 1982; Syková et al., 1974; Somjen et al., 1976; Heinemann et al., 1990; Amzica et al., 2002; Ding et al., 2016; Rasmussen et al., 2019). However, this technique creates a dead space surrounding the electrode, and can only measure from a single point in space, hence providing poor spatial resolution. Instead, to test our hypothesis, an optical approach like two-photon microscopy with fine-scale synaptic resolution seems ideal. Genetically-encoded green fluorescent protein-based [K^+^]_o_ sensors exist (Wu et al., 2022), which could be combined with a red-shifted genetically-encoded voltage indicator (Beck and Gong, 2019) to simultaneously monitor [K^+^]_o_ dynamics and synaptic spine V_m_ tuning *in vivo*. We hope that experimentalists in the future will use these advanced techniques to probe the existence of local dendritic [K^+^]_o_ hotspots, as well as their functions and the cellular mechanisms regulating them such as astrocyte-mediated K^+^ uptake (Wang et al., 2012).

In conclusion, our study is a first step towards unraveling the role of extracellular K^+^ changes in dendritic integration. Future theoretical and experimental studies are needed to obtain a comprehensive understanding of how local activity-dependent ionic shifts contribute to information processing and computations in dendrites.

## Methods

### Dendritic segments with orientation-tuned synaptic spines

We defined a dendritic segment as a cylinder with length L = 10 µm and a radius R_1_ = 1 µm (Takahashi et al., 2012; Iacaruso et al., 2017; Scholl et al., 2017; Kerlin et al., 2019). Each segment was randomly populated with between 7 and 13 synapses, yielding spine densities comparable to neocortical tissue (Comer et al., 2020; Parker et al., 2020; Iascone et al., 2020), and the orientation preference of individual spines was dependent on the synaptic organization type to which they belonged, that is, the diverse or similar input tuning regime. Spines within segments with similar tuning were assigned an orientation preference using a half-circular normal distribution with σ = 15^◦^ (Wilson et al., 2016) and a mean target orientation arbitrarily set to 0^◦^, while spines within segments with diverse tuning were given orientation preferences using the uniform distribution. All spines, irrespective of spatial organization regime, had a tuning curve width given by a half-circular normal distribution with σ = 11^◦^ heuristically determined, and spine activity for each tuning curve spanned from 0 to 1. Importantly, for both types of input regimes, we could reproduce the spine orientation preference distributions measured previously in the visual cortex of ferrets (Wilson et al., 2016). Moreover, the orientation preferences of our diverse tuning regime resembled previous those previously recorded in the visual cortex of mice (Jia et al., 2010).

### Dendritic spine activity factor

We modeled the activation of spines within dendritic segments by assuming that all spines were activated by visual stimulation according to their orientation tuning curve. To assess the expected synaptic spine activity level within a segment, we calculated the mean activity of all spines within that segment. We then obtained an activity factor by comparing the expectation values from the expected spine activity distributions for the two types of dendritic segments. We sampled 10^4^ dendritic segments and used a Gaussian kernel density estimator to determine the probability density function p(x) for the expected spine activity distributions for each segment type. The expectation value was calculated as **E**(X) = ∑_i_ x_i_p_i_, where x was a linearly spaced set of values between 0 and 1. To obtain the activity factor tuning curve (**Figure 1F**), we repeated this procedure for all stimulus orientations in the interval θ ∈ [0 : 90^◦^].

### Extracellular space surrounding the dendrite segment

The exact size and shape of the extracellular space surrounding functionally-mapped short dendritic segments is currently unknown, so we chose to model it as a cylinder that encapsulates the dendritic segment cylinder. The space in between these two cylinders thus constitutes the extracellular space in our investigations, and its volume is given by:

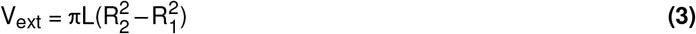

With R_1_ and R_2_ being the radius of the inner and outer cylinder, respectively, both with length L = 10 µm. This can be rewritten to express the volume ratio between the two cylinders:

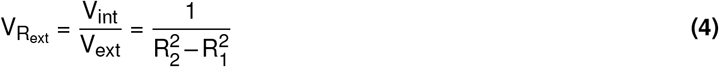

For simplicity, synaptic spines were considered points on the surface of the inner cylinder, and hence spine head volume was part of the dendritic segment volume.

### K^+^ diffusion state

During synaptic excitatory transmission, K^+^ ions are released into the extracellular space through primarily NMDA receptors (Rice and Nicholson, 1990; Shih et al., 2013; Poolos et al., 1987) and move around due to diffusion, although their movement is hindered by various cellular elements within the extracellular space. To account for this hindrance, we define effective diffusion as a function of tortuosity (Nicholson, 1993):

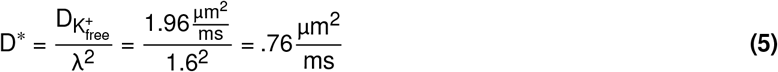

Here 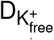 is the free ionic diffusion of a K^+^ particle in physiological saline, λ is a non-dimensional measure of how hindered an ion is in its movement in a given space, and D^∗^ is the resulting diffusion rate. The parameter values for 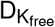 and λ were based on previous work (Nicholson, 1993; Halnes et al., 2016, 2013; Chen and Nicholson, 2000). Here we assume that diffusion in the radial direction is negligible, as the distance between the two cylinders is very small. Moreover, we assume closed boundary conditions on the outer cylinder wall, since the extracellular space here mainly contacts cellular membranes of neighboring neurons and/or glial cells. After a characteristic time of 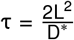, we assume that the [K^+^]_o_ is well-mixed and uniform throughout the outer cylinder. Using the dendritic segment length as the characteristic length, L = 10 µm, the characteristic time becomes τ ≈ 258 ms. Assuming that the majority of K^+^ efflux occurs within ≈ 100 ms from initial synaptic activation, we predict a peak [K^+^]_o_ elevation after ≈ 300 ms. Finally, we assume that the boundary between the dendritic segments is closed, implying a finite ion concentration within the segment. If K^+^ ions was not confined to one segment, we would expect ions on average to move ≈ 30 µm during the characteristic time before being well-mixed. This limits our approach since we do not investigate the effect of K^+^ ions moving along the dendrite, potentially affecting neighboring segments. However, given the average distance traveled for a K^+^ ion during the time period considered here, only the directly neighboring segment could potentially be affected, thus keeping the ionic shift very localized.

### K^+^ flux over the neuronal membrane

To estimate synaptic activity-dependent [K^+^]_o_ changes near dendritic segments, we assumed that local increases in [K^+^]_o_ are linearly correlated with the expected spine activity, as a result of K^+^ efflux from the intracellular space. By scaling the K^+^ flux in the diversely-tuned input regime with the orientation-dependent spine activity factor, we obtained an estimate of the K^+^ flux in the similarly-tuned regime. Using this, in conjunction with equation 4, we derived equations for the two types of input regimes as:

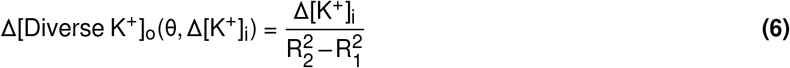

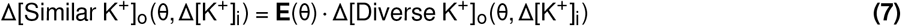

With Δ[K^+^]_o_ and Δ[K^+^]_i_ being the change in extracellular or intracellular [K^+^], respectively, R_1_ and R_2_ being the radius of the inner and outer cylinder, respectively, and **E** being the orientation-dependent spine activity factor. Using the Goldman-Hodgkin-Katz equation, we can express the change in 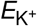 as:

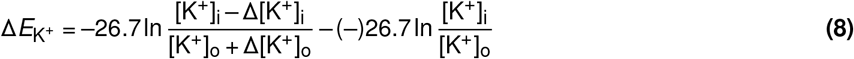

### Point-dendrite model

We simulated the dendritic segment V_m_ as an isolated resistance-capacitance circuit, similar to the common practice for point-neuron models (Dayan and Abbott, 2001). The major differences between point-neuron models and the point-dendrite model developed here, stems from the specific ion channel setup and cellular resistance, altered to mimic dendritic properties (**Table S1**). The circuit is given by the ordinary differential equation:

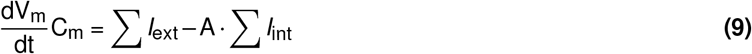

C_m_ is the membrane capacitance 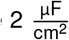, adjusted to recreate the contribution from the spine heads (Ujfalussy and Makara, 2020), *I*_ext_ is the current generated by extrinsic ion channels (AMPA and NMDA receptors), and *I*_int_ is the current generated by intrinsic ion channels (K_leak_, Na_V_, K_V_, K_M_, K_A_, K_Ca_, Ca_V_, and HCN). The latter is multiplied by the surface area, as these channels are scattered uniformly around the surface of the dendrite. All relevant channel dynamics and specific parameters are included in the Supplementary Materials. The system was simulated using an exponential Euler scheme (cnexp):

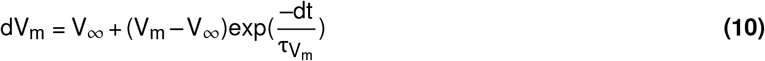

Here V_∞_ is the solution to the equation 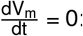:

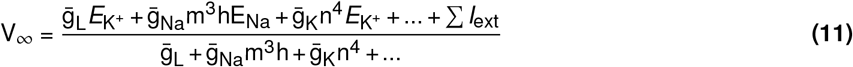

Here 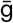 denotes a channel’s average conductance, n^x^ is from the individual channel dynamics controlling the gating and channel-specific conductance, and *E*_ion_ denotes the relevant reversal potential. The membrane time constant is given as:

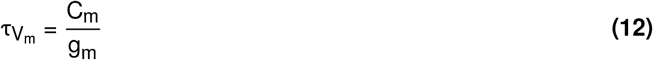

Here C_m_ is the capacitance of the membrane and g_m_ is the total membrane conductance.

The point-dendrite model was populated with orientation-tuned synaptic spines as described in the previous section. For each excitatory input stimulation burst, all spines were activated with a Poisson-distributed delay (λ = 30 ms). The orientation tuning of the individual spines was achieved by scaling the AMPA and NMDA currents with the spine’s tuning curve. To create a stimulation train consisting of several input bursts, we replicated the initial activation two times with a constant delay of 500 ms in between bursts. As we do not know the specific speed and kinetics of the activity-dependent local increase in [K^+^]_o_, we modelled it is a step function in between bursts: The first burst was regarded as the control condition with 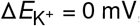 = 0 mV, and the two subsequent bursts as the experimental condition for a given 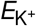 shift in the interval 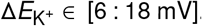 ∈ [6 : 18 mV]. We used a V_m_ of –30mV as a threshold criteria for identifying dendritic spikes. This value was chosen because when plotting the dendritic V_m_ as a function of synaptic input strength, we observed the largest change in V_m_ around this value, as synaptic integration transitions from subthreshold input summation to suprathreshold spiking.

### Linear dendritic spike function

The point-dendrite model was here modified so that all spines had a constant mean activity level. This modification allowed us to investigate the relation between mean spine activity (w; i.e., mean activity for the dendritic segment), 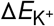, and the number of spines (N) more effectively. By varying each variable in small increments, we identified the lowest value of 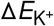 that was able to push the V_m_ over the threshold of –30mV, our working criteria for dendritic spike generation, given N and w. We assumed that the relation between the three parameters was linear, and we therefore fitted a plane to obtain a model of 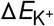 needed to trigger dendritic spikes as a function of N and w. For this we used the planar equation αw + βN + ν = 0 and solved it as Ax = b (**Figure S1**):

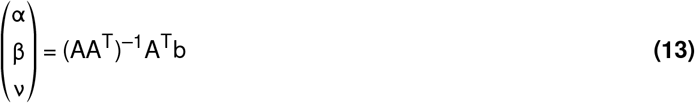

From this we could then estimate the dendritic spike probability as a function of 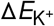 and stimulus orientation. Using the formalism for the setup of the dendritic segments with similarly-tuned inputs we sampled 10^4^ segments at each orientation and using the sampled expected spine activity and number of spines, we determined the 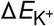 needed to generate a dendritic spike according to 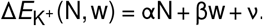 (N, w) = αN + βw + ν. To obtain 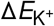 for a given orientation, we used a normalized version of the spine activity factor seen in **Figure 1F**, and multiplied the corresponding activity factor with the maximal 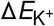 at the target orientation. Using this, we calculated the fraction of dendritic segments, at each orientation, that showed dendritic spikes for a given 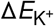 and converted this into a probability.

### Current-voltage attractor landscape

To understand the biophysical principles governing the regulation of dendritic spikes by local K^+^ changes we used dynamical systems theory (Strogatz, 2015). While our point-dendrite model is defined by a differential equation 9, it is not possible to solve it analytically. Rather, we chose to plot the exact solutions to the differential equations evaluated at different V_m_ as a vector plot, which in essence is similar to summing all I-V curves from the included intrinsic and extrinsic ion channels. On the first axis we thus have V_m_ and on the second we have the derivative which in our case can be denoted as 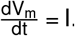. The time-dependency of the extrinsic ion channels was modelled in the same manner as for the point-dendrite model (see Supplementary Materials). Extrinsic ion channels were simulated in the time interval [0 : 100 ms] using the Euler method. For each time step, an I-V curve was generated and the resulting I-V curves were concatenated to create a time-dependent I-V landscape (**Figure 3F** and **3G**; **Movie S1**). To compare the effect of [K^+^]_o_ changes, we simulated the system with 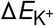 = 0 mV and 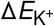 = 18 mV.

### Neuron model

Using the NEURON simulation environment as a backbone, we constructed a neuron model to test the role of local [K^+^]_o_ elevations on dendritic input integration and somatic output firing. The dendrite’s morphology was constructed as a fractal tree: for each dendritic section generation, three new sections, each with half the length of the former, was added. This gave a fractal with dimension D = ln 3/ ln 2 = 1.58, mimicking the generic compartmentalization and fractal dimensionality of the apical tree of pyramidal neurons (Smith et al., 2021; Porter et al., 1991). The fractal apical dendrite tree was extended with a trunk piece and a soma to complete the neuron morphology, and intrinsic ion channels (K_leak_, Na_P_, Na_T_, K_P_, K_T_, K_DR_, K_Ca_, K_M_, Ca_LVA_, Ca_HVA_, and HCN) were inserted into the cellular compartments (**Table S2**).

To investigate the compartmentalization of dendritic integration, we stimulated synaptic spines residing in segments with similar orientation tuning on the most extreme apical branches of the dendritic tree. For this, a Poisson-distributed number of dendritic segments (λ = 32 segments) were selected to host the inputs in each simulation iteration, and the number of spines and their orientation tuning were sampled as described above. The orientation tuning of individual spines was achieved by scaling the AMPA and NMDA currents with the spine’s tuning curve. All 10 µm dendritic segments within the fractal tree underwent 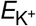 shifts depending on their type; that is, the diversely- or similarly-tuned type (1:1 ratio of types expect in **Figure S3**). The type of segment determined the magnitude of local 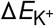 imposed during simulations, grouped into three conditions: no change (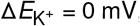 = 0 mV for both the diversely- and similarly-tuned type), small changes (Peak 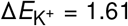 = 1.61 and 6 mV for the diversely- and similarly-tuned type, respectively), or large changes (Peak 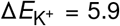 = 5.9 and 18 mV for the diversely- and similarly-tuned type, respectively), based on the [K^+^]_o_ changes found in **Figure 1**. To simulate a realistic somatic firing pattern, the spines within each dendritic segment were activated with a segment-relative random delay in the interval of 0 to 300 ms, and repeated three times with a constant 500 ms delay between stimulation burst. Each individual neuron setup was simulated with each of the three 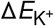 conditions to directly asses the effect of the local K^+^ changes.

### Neuronal activity measures

To obtain a better understanding of how the local [K^+^]_o_ changes impacted neuronal activity in the neuronal model we, in addition to the conventional measurement of somatic firing rate, introduced two additional measures: Neuronal firing gain transformation and area under the curve (AUC) for the integrated dendritic V_m_ signal.

#### Neuronal firing gain transformation

We assumed that the gain modulation of the somatic firing tuning curve could be described by a multiplicative or additive function, similar to previously described in the visual cortex of mice (Polack et al., 2013). The multiplicative and additive functions are given by:

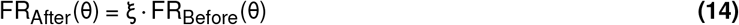

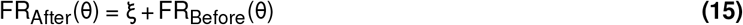

Here FR_Before_(θ) and FR_After_(θ) are the somatic firing rates at a given orientation with non-shifted or shifted 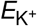 values, respectively, ξ is the fit parameter that describes the change in firing rate either as a gain coefficient or as a gain constant for the multiplicative and additive gain transformation, respectively. To determine the fit parameters, we fitted the two functions to the firing rate tuning curve data using the χ^2^ fitting method (**Table 1** and **Figure S2**). Only the multiplicative method was able to produce a satisfying fit and capture all features of the tuning curve, whereas the additive only captured the responses to orientations far away from the target orientation.

**Table 1.** Fitted gain parameters. Gain parameters obtained for the multiplicative (top) and additive functions (bottom) for the small and large 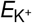 shifts (left and right, respectively). Errors were assumed Gaussian and reported as standard deviations. See also **Figure S2**.

| | $\xi$ Small $\Delta E_{K^+}$ | $\xi$ Large $\Delta E_{K^+}$ |
| --- | --- | --- |
| Multiplicative | $1.239 \pm 0.092$ | $1.964 \pm 0.107$ |
| Additive | $0.320 \pm 0.166$ | $0.348 \pm 0.102$ |

#### Area under the curve

To measure the strength of the integrated dendritic V_m_ signal, we computed the AUC of the V_m_ measured at the base of the dendritic trunk, over an interval of 375 ms to capture the full synaptic activation burst. To exclude the contribution of bAPs, we here turned off the voltage-gated sodium channels in the soma and trunk, which eliminated somatic firing. To calculate the AUC, we first subtracted the V_m_ baseline of the control signal, that is, 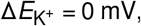 = 0 mV, from all traces, and used the trapezoidal method.

### Orientation selectivity index

The orientation selectivity index was computed as 1 – circular variance (CV) in orientation space given by:

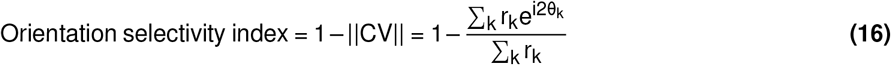

Where r_k_ is a measurable output at orientation θ_k_ (in radians); here we used dendritic spiking probability and somatic firing rate as output measures. To ensure a real-valued result, we took the modulus of CV: ||CV|| = 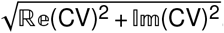.

## Statistics

To determine if somatic firing rates, obtained with either small or large local 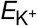 shifts, differed from firing rates obtained with no 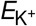 shift, we performed one-tailed Student’s *t* -tests on the residuals of the somatic firing rates. To determine if orientation selectivity indices, obtained with either no, small, or large 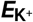 shifts, differed across conditions, we used a χ^2^ test to fit a zero-slope constant function to the indices from all three conditions. *P* < 0.05 was considered statistically significant, where \**P* < 0.05, \*\**P* < 0.01, and \*\*\**P* < 0.001.

## Data availability

All models used were implemented using Python version 3.10 and NEURON version 8.2. All models are available on Git: github.com/malthenielsen/potassium_hotspots or ModelDB (access number: 267732).

## Acknowledgments

We thank Akihiro Matsumoto, Alessandra Lucchetti, and Eva Maria Meier Carlsen for discussions and comments on the manuscript. R.N.R. and M.S.H. acknowledges support from the Lundbeck Foundation (R370-2021-764 and R347-2020-2250, respectively). M.H.J. and M.S.N. acknowledges support from the Independent Research Fund Denmark (9040-00116B) and the Novo Nordisk Foundation (NNF20OC0064978). A.P. acknowledges support from the Theodore Papazoglou FORTH Synergy Grants. N.T. acknowledges support from the University of Bordeaux Initiative of Excellence program, the Region Nouvelle-Aquitaine, the ATIP-Avenir program, and the Brain Science Foundation.

## Author Contribution

M.S.N., N.T., A.P., and R.N.R. conceived of the presented idea. M.S.N., N.T., A.P., M.S.H., M.H.J., and R.N.R. planned the project. M.S.N. developed the theoretical formalism, performed the analytic calculations and performed the numerical simulations. A.P., M.S.H., and M.H.J. verified the methodologies. M.S.N., N.T., A.P., and R.N.R. contributed to the interpretation of the results. M.S.N., N.T., A.P., and R.N.R wrote the manuscript with input from all authors.

