## Supplementary Material for "Local changes in potassium ions modulate dendritic integration"

##### Document inventory

1. Biophysical models
    - General setup
    - Intrinsic ion channels
    - Extrinsic ion channels
2. Tables
    - Point-dendrite model
    - Neuron model
3. Figures
    - Dendritic spike emergence plane fit
    - Gain function fit
    - Dendritic segment type ratio sensitivity
4. Movies
    - Animation of current-voltage attractor landscape during dendritic spike

### 1 Biophysical models

#### General setup

Below we describe the biophysical models used to create ion channel dynamics in the point-dendrite model. The models used for the neuron model in NEURON were very similar, but with some slight differences. For more information we refer to the enclosed .mod files found on Git: github.com/malthenielsen/potassium_hotspots or ModelDB (access number: 267732).

To mimic the physiological intra- and extracellular ionic environment we used the following concentrations:

- $[K^+]_i = 140\ mM \qquad [K^+]_o = 4\ mM \qquad (\Delta[K^+]_o \in [0.25 : 5\ mM])$
- $[Na^+]_i = 7\ mM \qquad [Na^+]_o = 140\ mM$
- $[Ca^{2+}]_i = 0.0001\ mM \qquad [Ca^{2+}]_o = 1.5\ mM \qquad$ (See $[Ca^{2+}]_i$ time dynamics below)

#### Intrinsic ion channels

Common for all intrinsic ion channels is that they conduct a single ion species ($Na^+$, $K^+$, or $Ca^{2+}$) and the reversal potential for that ion species is given by:

$$E_{ion} = \frac{RT}{z_{ion}F} \ln\left(\frac{[ion]_i}{[ion]_o}\right)$$

Here $R = 8.1415\frac{J}{mol;\ K}$ is the gas constant, $T = 311°K$ is the temperature in kelvin, $F = 96485.3321\frac{s\ A}{mol}$ is the Faraday constant, and $z_{ion}$ is the ion valence.

Some models also includes the $Q_{10}$ temperature-dependent coefficient denoted as $T_Q$:

$$T_Q = 2.3\frac{T-23}{10}$$

- **Leak K$^+$ channel (K$_{\mathbf{Leak}}$)**:

$$I_{K_{Leak}} = g_{Leak}(V_m - E_{K^+}) \tag{1}$$

- **Voltage-gated Na$^+$ channel (Na$_{\mathbf{V}}$)**:

$$m_\tau = \frac{m_\alpha + m_\beta}{T_Q} \qquad m_\infty = \frac{m_\alpha}{m_\alpha + m_\beta} \qquad \begin{cases} m_\alpha &= \frac{0.182(V_m+35)}{1-\exp-(V_m+35)/9.8} \\ m_\beta &= \frac{0.14(-V_m-35)}{1-\exp-(-V_m-35)/9.8} \end{cases}$$

$$h_\infty = \frac{1}{\frac{1+\exp V_m+65}{T_Q}} \qquad h_\tau = \frac{h_\alpha + h_\beta}{T_Q} \qquad \begin{cases} h_\alpha &= \frac{0.024(V_m-50)}{1-\exp-(V_m-50/5} \\ h_\beta &= \frac{0.0091(-V_m+75)}{1-\exp-(-V_m+75)/5} \end{cases}$$

$$\frac{dm}{dt} = \frac{m_\infty - m}{m_\tau} \qquad \frac{dh}{dt} = \frac{h_\infty - h}{h_\tau}$$

$$I_{Na_V} = T_Q g_{Na} m^3 h((V+25) - E_{Na^+})$$

- **Voltage-gated $K^+$ channel ($K_V$)**:

$$n_\tau = \frac{n_\alpha + n_\beta}{T_Q} \qquad n_\infty = \frac{n_\alpha}{n_\alpha + n_\beta} \qquad \begin{cases} n_\alpha & = 0.02\frac{V_m - 25}{1 - \exp\frac{-V_m - 25}{9}} \\ n_\beta & = -0.006\frac{V_m - 25}{1 - \exp\frac{V_m - 25}{9}} \end{cases}$$

$$\frac{dn}{dt} = \frac{n_\infty - n}{n_\tau}$$

$$I_{K_V^+} = T_Q g_{K^+} n(V_m - E_{K^+})$$

- **M-type $K^+$ channel ($K_M$)**:

$$n_\tau = \frac{n_\alpha + n_\beta}{T_Q} \qquad n_\infty = \frac{n_\alpha}{n_\alpha + n_\beta} \qquad \begin{cases} n_\alpha & = 0.001\frac{(V_m + 30)}{1 - \exp\frac{-V_m + 30}{9}} \\ n_\beta & = -0.001\frac{(V_m + 30)}{1 - \exp\frac{-V_m + 30}{9}} \end{cases}$$

$$\frac{dn}{dt} = \frac{n_\infty - n}{n_\tau}$$

$$I_{K_m^+} = T_Q g_{K^+m} n(V_m - E_{K^+})$$

- **A-type $K^+$ channel ($K_A$)**:

$$\zeta = \frac{-1.5}{1 + \frac{\exp V_m + 40}{5}}$$

$$n_\tau = \frac{n_\beta}{0.05 T_Q (1 + n_\alpha)} \qquad n_\infty = \frac{1}{1 + n_\alpha} \qquad \begin{cases} n_\alpha & = \exp\frac{96.48\zeta(V_m - 11)}{8.13(273 + 38)} \\ n_\beta & = \exp\frac{53.03\zeta(V_m - 11)}{8.13(273 + 38)} \end{cases}$$

$$l_\tau = .26(V_m - 50) \qquad l_\infty = \frac{1}{1 + l_\alpha} \qquad \begin{cases} l_\alpha & = \exp\frac{96.48\zeta(V_m - 11)}{8.13(273 + 38)} \end{cases}$$

$$\frac{dn}{dt} = \frac{n_\infty - n}{n_\tau} \qquad \frac{dl}{dt} = \frac{l_\infty - l}{l_\tau}$$

$$I_{K_A^+} = g_{K_A^+} nl(V_m - E_{K^+})$$

- **$Ca^{2+}$-gated $K^+$ channel ($K_{Ca}$)**:

$$n_\tau = \frac{n_\alpha + n_\beta}{T_Q} \qquad n_\infty = \frac{n_\alpha}{n_\alpha + n_\beta} \qquad \begin{cases} n_\alpha & = 0.01[Ca^{2+}]_i \\ n_\beta & = 0.02 \end{cases}$$

$$\frac{dn}{dt} = \frac{n_\infty - n}{n_\tau}$$

$$I_{K_{Ca}^+} = T_Q g_{K_{Ca}^+} n(V_m - E_{K^+})$$

- **Voltage-gated $\mathbf{Ca^{2+}}$ channel ($\mathbf{Ca_V}$)**:

$$m_\tau = \frac{1}{m_\alpha + m_\beta} \qquad m_\infty = \frac{m_\alpha}{m_\alpha + m_\beta} \qquad \begin{cases} m_\alpha &= 0.055 \frac{-27 - V_m}{\exp \frac{-27 - V_m}{3.8} - 1} \\ m_\beta &= .94 \exp \frac{-75 - V_m}{17} \end{cases}$$

$$h_\tau = \frac{1}{h_\alpha + h_\beta} \qquad h_\infty = \frac{h_\alpha}{h_\alpha + h_\beta} \qquad \begin{cases} h_\alpha &= 0.000457 \exp \frac{-13 - V_m}{50} \\ h_\beta &= \frac{0.0065}{\exp \frac{-V_m - 15}{28} + 1} \end{cases}$$

$$\frac{dm}{dt} = \frac{m_\infty - m}{m_\tau} \qquad \frac{dh}{dt} = \frac{h_\infty - h}{h_\tau}$$

$$I_{Ca_V} = g_{Ca_V} m^2 h (V_m - E_{Ca^{2+}})$$

The $[Ca^{2+}]_i$ and degradation was modeled as:

$$\dot{[Ca]}_i = -20(A \cdot I_{Ca_V} + I_{NMDA}) - \frac{[Ca^{2+}]_i}{121.4} + \mathcal{N}(0, 3 \cdot 10^{-7})$$

Here A is the surface area of the dendrite and $I_{channel}$ is the current passing at a given time. $\mathcal{N}(\mu, \sigma)$ is a random number from a normal distribution with mean $\mu$ and standard deviation $\sigma$.

- **Hyperpolarization-activated cyclic nucleotide–gated channels (HCN):**
This is the only intrinsic ion channel with mixed ion conductance, that is, both $Na^+$ and $K^+$ moves through it. We can calculate the combined reversal potential as:

$$E_{HCN} = \frac{RT}{z_{ion}} \ln \left( \frac{p_{Na}[Na^+]_i + p_K[K^+]_i}{p_{Na}[Na^+]_o + p_K[K^+]_o} \right)$$

Here we assume that $p_{Na} = p_K$

$$m_\tau = \frac{1}{m_\alpha + m_\beta} \qquad m_\infty = \frac{m_\alpha}{m_\alpha + m_\beta} \qquad \begin{cases} m_\alpha &= \frac{0.00643(V_m + 154.9)}{\exp \frac{V_m + 154.9}{11.9} - 1} \\ m_\beta &= 0.193 \exp \frac{V_m}{33.1} \end{cases}$$

$$\frac{dm}{dt} = \frac{m_\infty - m}{m_\tau}$$

$$I_{HCN} = g_{HCN} m (V_m - E_{HCN})$$

#### Extrinsic ion channels

The extrinsic ion channels conduct several ion species and the reversal potential is given by:

$$E_{channel} = \frac{RT}{z_{ion}} \ln \left( \frac{p_{Na}[Na^+]_i + p_K[K^+]_i + p_{ca}[Ca^{2+}]_i}{p_{Na}[Na^+]_o + p_K[K^+]_o + p_{ca}[Ca^{2+}]_o} \right)$$

Here we assume that $p_{Na} = p_K = p_{Ca}$

- **AMPA receptor**:

$$\frac{dA}{dt} = \frac{-A}{\tau_1} \quad \tau_1 = 0.5ms \qquad \frac{dB}{dt} = \frac{-B}{\tau_2} \quad \tau_2 = 1.5ms$$

$$I_{AMPA} = g_{AMPA}(B - A)(V_m - E_{AMPA}) \quad \begin{cases} g_{AMPA} &= 5 \cdot 10^{-8} \end{cases}$$

- **NMDA receptor**:

$$Mg_{block}(V_m) = \frac{-1}{1 + [Mg^{2+}]_o/3.57exp(-0.1V_m)}$$

$$\frac{dA}{dt} = \frac{-A}{\tau_1} \quad \tau_1 = 4ms \qquad \frac{dB}{dt} = \frac{-B}{\tau_2} \quad \tau_2 = 42ms$$

$$I_{NMDA} = g_{NMDA}(B - A)Mg_{block}(V_m)(V_m - E_{NMDA}) \quad \begin{cases} g_{NMDA} & = -2.9{\cdot}10^{-7} \\ [Mg^{2+}]_o & = 2mM \end{cases}$$

### 2 Tables

| Channel | Conductance [$\mu Scm^{-2}$] |
|---|---|
| $g_{Leak}$ | 0.005 |
| $g_{Na_V}$ | 5.5 |
| $g_{K_V}$ | .2 |
| $g_{K_M}$ | 0.1 |
| $g_{K_A}$ | .9 |
| $g_{K_{Ca}}$ | .06 |
| $g_{Ca_V}$ | .12 |
| $g_{HCN}$ | .01 |

**Table S1.** Active conductances of the point-dendrite model.

| Channel | Channel (NEURON) | Dendrite [$Scm^{-2}$] | Trunk [$Scm^{-2}$] | Soma [$Scm^{-2}$] |
|---|---|---|---|---|
| $g_{Leak}$ | *pas* | $6{\cdot}10^{-5}$ | $6{\cdot}10^{-4}$ | $3{\cdot}10^{-6}$ |
| $g_{Na_P}$ | $g_{Nap}$ | - | - | 0.00172 |
| $g_{Na_T}$ | $g_{NaTa}$ | 0.021489 | 0.021489 | 2.04 |
| $g_{K_P}$ | $g_{K_per}$ | - | - | 0.0023 |
| $g_{K_T}$ | $g_{K_tst}$ | - | - | 0.0821 |
| $g_{K_{DR}}$ | $g_{SK+v3_1}$ | $18.08{\cdot}10^{-4}$ | $18.08{\cdot}10^{-4}$ | 0.438029 |
| $g_{K_M}$ | $g_{I_m}$ | $9.9{\cdot}10^{-4}$ | $9.9{\cdot}10^{-4}$ | - |
| $g_{K_{Ca}}$ | $g_{SK_{E2}}$ | $.02{\cdot}10^{-4}$ | $.02{\cdot}10^{-4}$ | $9.65{\cdot}10^{-2}$ |
| $g_{Ca_{LVA}}$ | $g_{Ca_{LVA}}$ | $22.7{\cdot}10^{-4}$ | $22.7{\cdot}10^{-4}$ | $3.34{\cdot}10^{-4}$ |
| $g_{Ca_{HVA}}$ | $g_{Ca_{HVA}}$ | $7.01{\cdot}10^{-4}$ | $7.01{\cdot}10^{-4}$ | - |
| $g_{HCN}$ | $g_{Ih}$ | $1.5{\cdot}10^{-5}$ | $1.5{\cdot}10^{-5}$ | $7.5{\cdot}10^{-5}$ |

**Table S2.** Active conductances of the neuron model.

### 3 Figures

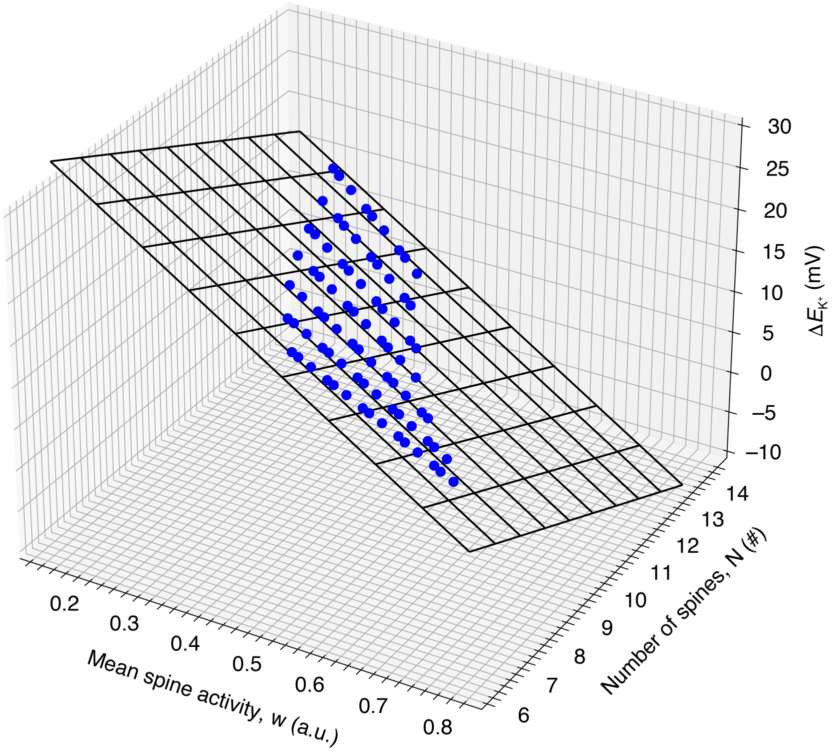


**Figure S1. Dendritic spike emergence plane fit.**
We created multiple input-output curves by varying either the mean spine activity (w), number of spines on dendritic segment (N), or $\Delta E_{K+}$, while keeping the two other input parameters constant, and having dendritic spike occurrence ($V_m$ above $-30$ $mV$) as the output measure. This was done multiple times for each condition and the resulting scatter of points in 3D space is shown here (blue circles). Moreover, we fitted a plane to this data (black grid), as we assumed the relation between w, N, $\Delta E_{K+}$ and dendritic spike generation was linear; points on the plane denote the parameter sets which were able to generate a dendritic spike.

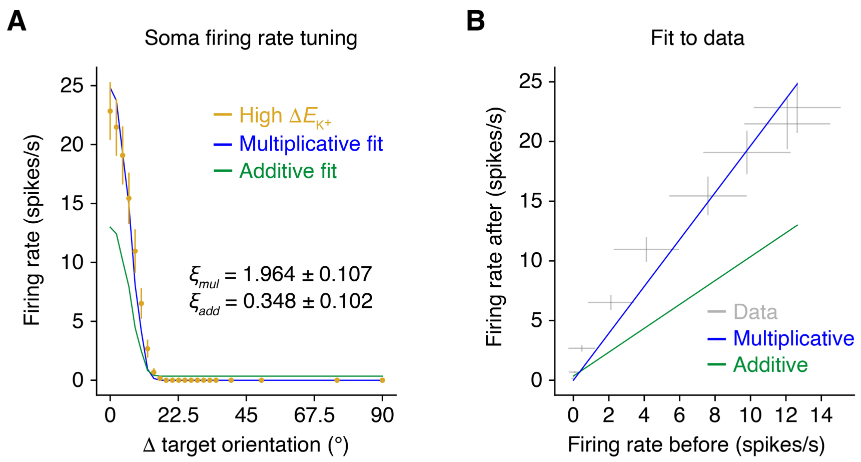


**Figure S2. Gain function fit.**
(**A**) Multiplicative and additive gain function predictions plotted together with the original soma firing rate data for the high $\Delta E_{K+}$ condition as a function of stimulus orientation relative to target orientation. The $\xi$ values denote the fit parameter that describes the firing rate change either as a gain coefficient or as a gain constant for the multiplicative ($\xi_{mul}$) and additive ($\xi_{add}$) gain transformation, respectively. Predictions were generated by either adding $\xi_{add}$ or multiplying $\xi_{mul}$ to the tuning curve with no $E_{K+}$ change (curve not shown). Errors were assumed Gaussian and reported as $\pm$ standard deviations.
(**B**) Actual multiplicative and additive gain function fits plotted together with the original soma firing rate data with no $E_{K+}$ change (firing rate before) against high $E_{K+}$ change (firing rate after). The original firing rate data was plotted as $\pm$ standard deviations on each axis. To fit the data we have only considered data with non-zero standard deviation.

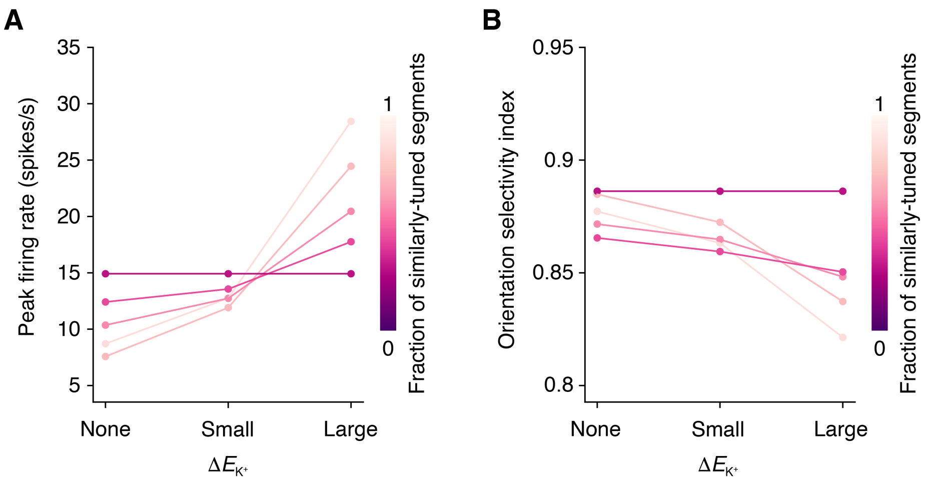


**Figure S3. Dendritic segment type ratio sensitivity.**
(**A**) Soma peak firing rate at target orientation as a function of dendritic $E_{K+}$ shift magnitude and fraction of dendritic segments with similarly-tuned inputs in the dendritic tree.
(**B**) Orientation selectivity index as a function of dendritic $E_{K+}$ shift magnitude and fraction of dendritic segments with similarly-tuned inputs in the dendritic tree.

### 4 Movies

**Movie S1. Animation of current-voltage attractor landscape during dendritic spike.**
(**A**) I-V curves including intrinsic ion channels and NMDA receptors during the generation of a dendritic NMDA spike without and with $E_{K^+}$ shift. Solid and open points indicate stable and unstable fixed points, respectively, and arrows indicate system flow direction.
(**B**) Simplified outline of the approximate $V_m$ during the dendritic NMDA spike with and without $E_{K^+}$ shift, highlighting the voltage levels the $V_m$ is attracted toward in **A**.
(**C**) Heatmaps showing the temporal evolution of the I-V curves without and with $E_{K^+}$ shift. The red line indicates the corresponding I-V curves shown in **A**, full and dotted white lines indicate stable and unstable fixed points, respectively, and the magenta and green lines being drawn indicate the instantaneous $V_m$ for the system without and with $E_{K^+}$ shift, respectively.
